# From 2D to 4D: a Containerized Workflow and Browser to Explore Dynamic Chromatin Architecture

**DOI:** 10.1101/2025.07.13.664622

**Authors:** David H. Rogers, Cullen Roth, Cameron Tauxe, Jeannie T. Lee, Christina R. Steadman, Karissa Y. Sanbonmatsu, Anna Lappala, Shawn R. Starkenburg

**Affiliations:** Information Sciences, Los Alamos National Laboratory, Los Alamos, NM, US; Genomics & Bioanalytics Group, Los Alamos National Laboratory, Los Alamos, NM, US.; Neptune, Inc. & Vistar Media, New York City, NY, US.; Department of Molecular Biology, Massachusetts General Hospital, Boston, MA, US.; Department of Genetics, Harvard Medical School, Boston, MA, US.; Theoretical Biology & Biophysics Group, Los Alamos National Laboratory, Los Alamos, NM, US.; New Mexico Consortium, Los Alamos, NM, US.

**Keywords:** 3D Visualization, Genome Modeling, Epigenomics, Hi-C, chromatin architecture, 3D Genome Browser

## Abstract

**Background:** Characterizing the physical organization of the genome is essential for understanding long-range gene regulation, chromatin compartmentalization, and epigenetic accessibility. Hi-C experiments generate two-dimensional (2D) genome-wide contact maps of chromatin interactions by capturing the spatial proximity between genomic loci, which reveal interaction frequencies but lack the spatial resolution needed to interpret the three-dimensional (3D) genome structure(s). Emerging evidence suggests that epigenetic regulation is closely linked to 3D genome architecture, and that structural changes over time (4D) drive key biological processes in development, disease, and environmental response. Thus, integrating 3D structure with functional data is critical for a more complete understanding of genome regulation. Previous work, most notably the 4DHiC chromosome modeling framework, has shown that physical multi-dimensional modeling approaches rooted in polymer physics and molecular dynamics can resolve these structures at biologically meaningful resolutions by integrating temporal Hi-C data with physical constraints to uncover dynamic chromosome reorganization. Thus, molecular dynamics simulations, constrained by Hi-C contact matrices, can resolve fine-scale structural changes and reveal functionally significant transitions in chromatin conformation.

**Results:** Herein, we present the 4D Genome Browser Workflow (4DGBWorkflow) and the 4D Genome Browser (4DGB). The algorithm is based on the 4DHiC method and the containerized tool is an end-to-end workflow that can transform, filter, and view 4D epigenomics and chromatin datasets, allowing non-specialists to apply three-dimensional modeling principles to diverse datasets and experimental conditions. The software executes on a laptop running macOS, Linux or Windows. From input Hi-C files (.hic), the 4DGBWorkflow produces 3D reconstructions of chromosomes, integrates the reconstruction with track data (e.g., epigenetic marks, transcriptome profiles), and provides comparative visualization of the results in a single workflow.

**Conclusions:** The 4DGBWorkflow and 4D Genome Browser are open-source tools for comparative analysis and visualization of 4D chromosome datasets, including chromatin architecture and epigenomic signals. Automatic integration of Hi-C data with molecular dynamics democratizes the construction of time resolved 3D genome structures, simplifying complex simulations and data integration schemes.

## Background

Understanding the three-dimensional (3D) architecture of the genome has become increasingly critical for deciphering the regulatory mechanisms underlying gene expression, epigenetic states, and cellular identity [2–4]. Beyond the linear sequence, the physical conformation of chromatin within the nucleus influences enhancer-promoter contacts, replication timing, and compartmentalization of active versus inactive domains [5–7]. Hi-C experiments, and related methods, have demonstrated that genomic loci interact over large distances in the nucleus, forming reproducible structures such as chromatin loops, topologically associating domains (TADs), compartments, and whole-chromosome territories [8–10]. These structural features are not static: they change during development, cell cycle progression, and in response to environmental stimuli or disease states [4, 11–14]. For example, the folding and unfolding of chromatin can determine whether genes are accessible for transcription, while epigenetic modifications such as histone acetylation or DNA methylation can further refine gene activity [15, 16]. Thus, the genome is not merely a static blueprint but a responsive, adaptable architecture whose regulation occurs at multiple spatial and temporal scales.

Yet, despite these insights, there remains a substantial gap in our ability to connect genome structure to function. High-resolution, genome-wide modeling of chromatin architecture that integrates structural, epigenetic, and transcriptional information is still a major methodological and computational challenge [17]. Traditional imaging approaches, such as fluorescence in situ hybridization, super-resolution microscopy, and cryo-electron tomography, have yielded critical visual confirmation of chromatin organization but are limited in resolution and throughput [18]. They cannot, for example, pinpoint the spatial relationships of specific regulatory elements or capture the broader organizational logic of the genome across multiple conditions or cell types.

By capturing the frequency of physical contacts between genomic regions, Hi-C provides the data necessary to reconstruct 3D models of entire genomes [19]. These models enable researchers to examine the spatial arrangement of chromosomes, identify changes in domain-level organization, and study how genome folding influences gene regulation [20, 21]. However, Hi-C data alone do not convey chromatin state—they must be complemented with data from assays such as ATAC-seq (chromatin accessibility), ChIP-seq (histone marks and transcription factors), and DNA methylation profiling to contextualize the structural information [12]. Without these layers, 3D models are descriptive but incomplete.

Even when multi-omics datasets are available, the computational burden of processing and integrating them is substantial. Constructing accurate 3D models from Hi-C data involves complex simulation algorithms, optimization of contact distance matrices, and parameter tuning [12]. Many existing tools require users to navigate multiple software environments, manage dependencies, and operate on high-performance computing systems or cloud-based infrastructures. This imposes significant barriers for non-specialist users, restricts reproducibility, and often leads to inconsistent or inaccessible results.

To overcome these challenges and democratize access to 3D genome modeling, we have developed the 4D Genome Browser Workflow (4DGBWorkflow) and Browser (4DGB, Figure 1), a standalone, containerized computational toolkit that enables researchers to construct, visualize, and compare high-resolution 3D genome models directly from their own data. This pipeline builds directly upon the 4DHiC framework introduced by Lappala et al. [12], which integrates Hi-C contact data with molecular dynamics simulations to generate time-resolved, physically grounded 3D genome reconstructions. The 4DGBWorkflow extends this framework into a reproducible and user-friendly software system. It supports the inclusion of other epigenomic signals, such as ATAC-seq, ChIP-seq, and RNA-seq data, which can be projected onto the 3D genomic models. This data integration allows users to overlay chromatin accessibility and regulatory features onto 3D models for a comprehensive view of structure-function relationships. In this work, we demonstrate the utility of the 4DGB software across use cases including developmental time courses and perturbation experiments. By enabling scalable, reproducible, and accessible 4D genome modeling, this tool represents a critical step forward in decoding the spatial and functional relationships of the genome.

**Figure 1:**
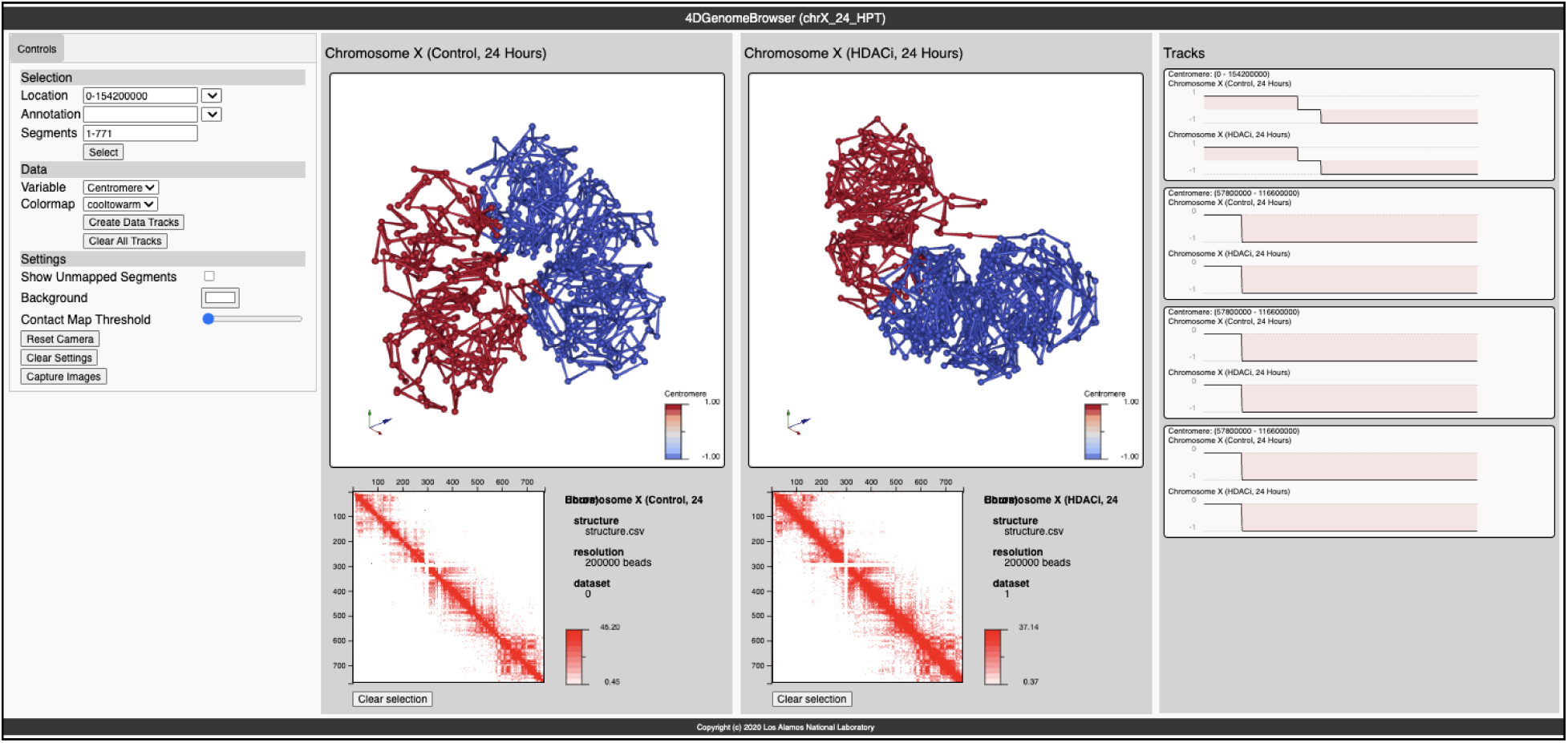
Screen capture of the 4D Genome Browser, demonstrating the fusion of track data with 3D structure(s). Center: Images represent modeling in 3D of chromosome X from A549 cells. Models were generated from Hi-C data gathered from control (left) and HDACi (histone deacetylase inhibitor) treated (right) samples, post 24 hours. Visualization in 3D depicts compactions of p and q chromosome arms (red and blue colors, respectively). Bottom: Hi-C data from control and HDACi treated maps, left and right, respectively [1]. Right: Data tracks showing centromere values along chromosome X. Along chromosome X, the p and q arms were given a binary coding of 1 and -1 (respectively) with the centromere locations begin assigned a value of zero. These values were mapped to the 3D renderings (middle) to display p and q chromosome arms in 3D space.

## Implementation

The 4DGB toolkit is an application that makes a complex workflow available to scientists as a cross-platform command line tool (Figure 2). Designed for ease of use and installation, the 4DGB software suite is implemented within a Docker container [22] enabling one-step deployment on target platforms. It encapsulates all necessary software components, eliminating complex installations and ensuring consistency across users and systems. The workflow includes modules for data preprocessing, contact map generation, 3D structural modeling via molecular dynamics simulations, annotation with functional tracks, and interactive visualization. Unlike current platforms that rely on centralized, cloud-based servers with limited flexibility, this tool empowers researchers to maintain full control over their data while facilitating rapid hypothesis generation and validation. The models generated satisfy a high proportion of experimental contact constraints (*>* 90%) and can be visualized through a local or web-based interface. In doing so, the 4D Genome Browser bridges the gap between raw experimental output and actionable biological insight, providing a framework to explore genome architecture as a functional entity, not merely a structural scaffold.

**Figure 2:**
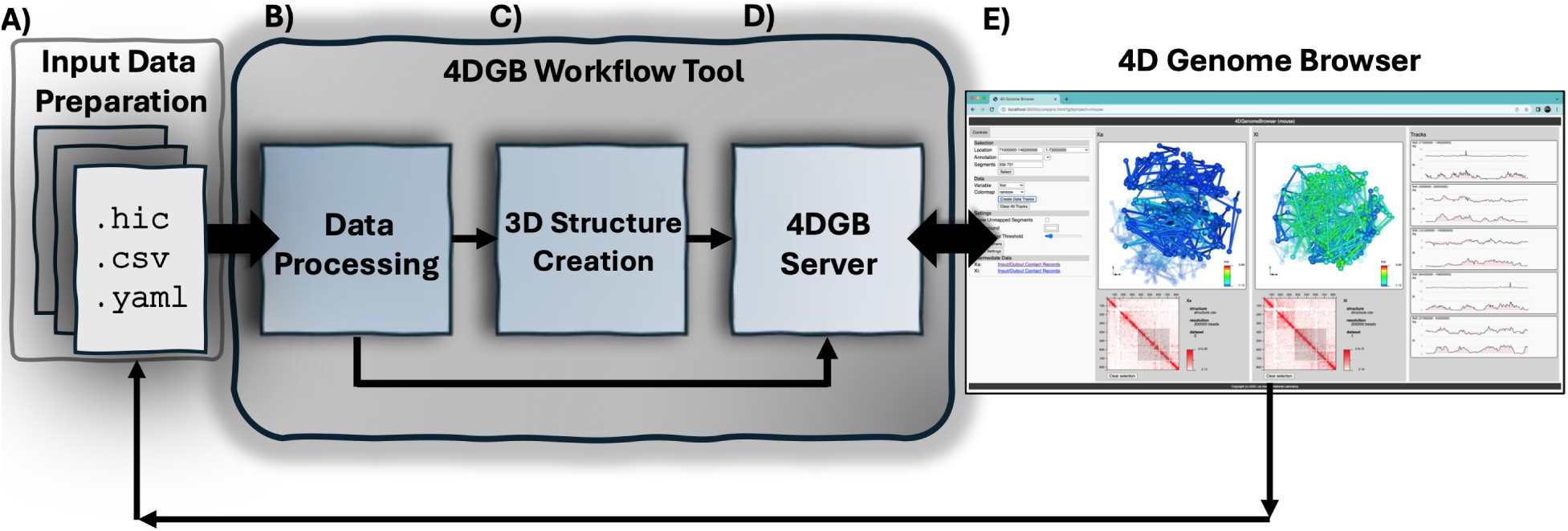
Diagram of the workflow described in this paper, which uses the 4D Genome Browser Workflow tool. There are three main steps in using the workflow: the user (A) defines a set of input data consisting of 1D and 2D data, (B–D) runs the workflow, and (E) then views the results in a web browser with the 4D Genome Browser. The 4D Genome Browser Workflow tool performs three steps: (B) Data processing and data fusion of input datasets to products needed by the workflow, (C) 3D structure creation using a molecular dynamics simulation (LAMMPS), and (D) a data server that provides (E) a novel browser view for comparative visualization of the results. After viewing the results, the user can add, delete or update input data to change the visualization. For example, more tracks can be added and visualized in the browser. This complex workflow is packaged in a container that can be easily installed and run on any desktop or laptop that supports Docker.

There are three high level steps in using the tool: preparing input data, running the workflow, and visualizing the results. The tool can be run iteratively on a collection of data, so that new tracks and features can be added over time. Complex operations, including the generation of 3D structural models, are executed conditionally, ensuring they are run only when the input data is new or updated. The tool can be used on common-sized data sets (on the order of mammalian chromosomes including mouse and human) and run on a laptop. The workflow and browser can also be run on a web-based system (discussed below) with access to more compute power. The 4DGB tool is designed for ease of implementation (from an example project) and ease of use. Detailed instructions on the latest released version of the tool can be found in the online documentation.

A significant contribution of the 4DGB tool is that it builds upon the 4DHiC modeling framework [12] to transform 1D and 2D input data—such as .csv and .hic files—into high-resolution 3D genome models. The 4DGB pipeline performs all necessary operations to generate 3D chromatin geometry, integrate signals from associated omics data (such as ATAC-seq and ChIP-seq), and produce a comparative visualization of the results. Using principles from polymer physics and molecular dynamics, 4DHiC applies Hi-C–derived harmonic constraints to drive the folding of a self-avoiding polymer chain into biologically plausible chromosome conformations. The 4DGBWorkflow automates this process in a containerized environment: the user simply defines a set of input files, and the workflow handles internal data preparation, execution of 4DHiC-based simulations, and downstream data fusion for interactive visualization. [12, 23, 24].

The 4DGB tool can be installed by the user with a single command. The installed tool then manages the download of the Docker images as needed to complete operations requested by the user. Docker was chosen among alternative containerization solutions, such as Podman [25] or CharlieCloud [26], for its ubiquity in the industry. Furthermore, Docker provides a simplified install process, especially for non-Linux platforms compared to the alternatives.

Before running the tool, the user must prepare input data (Figure 2A). A combination of .hic and .csv files define the datasets that the user would like to compare in the final visualizations. The tool and workflow is expecting:

1. Contact Maps. Two comparable Hi-C data files (.hic), e.g., a control file and a condition file containing the same set of named chromosomes. This data is required.
2. Track Data. One or more sets of track data to be visualized on the structure that is generated by the workflow. This information must be in the .csv format. These track files are optional, and the workflow will run without them.
3. Project Information. A project.yaml file that defines metadata and organization for the input datasets. This file can be very simple, but there are options to include different kinds of data. An example of a complete project.yaml is included with the workflow template, and examples can be found in the online documentation. There are several sections defining that data with vocabularies as follows:

(a) workflow: This is metadata about the workflow.
(b) project: This section defines information and settings for the project.
(c) datasets: This section defines .hic files for the workflow.
(d) tracks: This defines track data that can be painted on the 3D structures created.
(e) annotations: This defines annotation files that can be used to select regions in the 4D Genome Browser. Either .gff or .csv files can be used.
(f) bookmarks: This section defines features and locations of interest that can be quickly selected in the 4D Genome Browser.

Raw Hi-C data (in the format of paired-end .fastq files) can be processed into .hic format using pipelines such as Juicer [27] or SLUR(M)-py [28], which are compatible with the 4DGBWorkflow. The downstream protocol—from Hi-C data preprocessing, binarization, and matrix thresholding, to molecular dynamics simulation and 3D structure generation—follows the 4DHiC framework [12]. This approach introduced a rigorous and physically grounded methodology for transforming Hi-C contact maps into dynamic, time-resolved 3D chromosome reconstructions using LAMMPS. The 4DGBWorkflow and Browser containerize and streamline this process, offering a graphical interface for comparative visualization, with the scientific and computational core—including matrix handling, particle interaction logic, constraint implementation, and convergence validation derived from the 4DHiC protocol. Each input .hic file must contain contact records for a target chromosome and resolution (e.g., chr22 at 200 kb), and while multiple chromosomes may be present, the current implementation operates on a single chromosome-resolution pair per run.

As in the original 4DHiC implementation, the workhorse behind the preprocessing is the hic-straw python module from the Juicer suite of tools [27]. Each .hic file is parsed using hic-straw and Knight-Ruiz standardized contact counts are brought in for analysis [29]. Contacts along the diagonal of the Hi-C matrix, which represent linear, self-contacts within a chromosome (i.e., *x_i_* = *y_j_*, where x and y are the rows and column coordinates of the Hi-C matrix) are removed from analysis. The Hi-C matrix is then transformed into binary values using a count threshold (e.g., count threshold: 2.0), removing contact values between Hi-C bins lower than this threshold (setting to zero) and setting all others to one. Parameter values for Hi-C data filtering and processing (i.e. chromosome, resolution, and count threshold) are set within a project.yaml file, discussed in later sections and shown in the online documentation. This binary Hi-C contact map is then passed to the modeling algorithm, the Large-Scale Atomic/Molecular Massively Parallel Simulator (LAMMPS), for generating a 3D model of the chromosome [12, 24].

### Running The 4DGB Workflow Tool

After data preparation, a user can run the workflow (Figure 2C). This executes the data processing, computes 3D structures (using LAMMPS [12, 23, 24]), and runs a local web server on the resulting data. The data can then be viewed on a local browser. The implementation of these steps is hidden from the user, but behind the scenes, a python script manages the downloading and execution of docker images for these steps. Together, the script and docker containers manage a complex series of steps that transforms 1D and 2D input data into a 4D visualization of a single chromosome. This allows scientists to view track data in a 3D context, and to compare different experiments or states of the data to each other. The workflow can be run iteratively on a collection of data, such that tracks, annotations and other data can be added over time. The workflow will only re-run expensive steps, such as computation of 3D structure, when relevant input data has changed.

The 4DGB software operates on a collection of data known as a ‘project’, all contained within the same project directory. Output from the operations is written to a .build directory that is created within the project directory. The output is text files that can be viewed with common tools, and can be managed with ordinary shell file operations. There are three operations within the 4DGB workflow that may be performed on a set of project data; these are named run, build, and view.

**run**: This will first build, then view a project, with the constraints noted in the steps below. For example, it will only perform the build step if a valid 3D structure does not exist for the data.

**build**: This will take the project input data and create a 3D structure from the .hic files by running a LAMMPS molecular dynamics simulation. This is the most computationally expensive step of the workflow, and the tool will only rerun the build step later if it detects that the input .hic files and the computed 3D structure are out of sync.

**view**: This will run a 4D Genome Browser server on data from a completed build. Viewing a project that has not been built will not fail, but instead will result in a browser with no content.

After invoking the run command, the workflow manages several linked steps, each of which is described in more detail below.

**Data Processing** (Figure 2B). The user’s data is prepared for both inputs to the molecular dynamic simulation(s) and template files for the 4D Genome Browser Workflow. This includes preparing an input deck and initial conditions datasets for the simulation used in 3D structure creation, and organizing the tracks and other data called out in the project.yaml into data structures needed by the 4DGB server.

**3D Structure Creation** (Figure 2C). The 3D structure is created using LAMMPS and follows protocols established in earlier work [12, 24]. The initial configuration forms an open coil chain as a result of a self-avoiding random walk. Experimental data-driven Hi-C constraints were used directly such that connected pairs (virtual Hi-C bonds) were required to form harmonic bonds between interacting particles, and those were directed to form connected pairs prior to production simulations. Once over 80% of data-driven bonding constraints were satisfied, the convergence was measured by comparing the experimental contact maps to the simulated ones by computing the Pearson correlation coefficient. The result of this setup is a connected series of 3D positions that define a configuration of the chromosome that satisfies the constraints defined in the input .hic file.

Experimental Hi-C data contains unmappable regions where DNA sequences are undefined, unable to be sequenced, or lack mapped sequenced data, such as centromeres, and it is impossible to know the exact connectivity of certain segments [12]. In our model, the unmappable regions ‘loop out’ from the final folded state due to the fact that these regions have no specified Hi-C constraints. These regions can be viewed or hidden in the final visualization, and are typically excluded from any further analyses.

**Execution of 4DGB Server** (Figure 2D). Finally, an instance of the 4DGB server is started. This server first organizes the 3D structure and track datasets in a database and provides an API for a browser application to query. The server then waits to be connected to by a locally running browser (Figure 2E).

**Visualization of Data in Local Browser**. After the workflow has completed successfully, a URL is printed to the shell. The user can cut and paste that URL into a web browser to interactively view the datasets. This runs the 4D Genome Browser for comparative visualization of the results. Figure 3 shows the browser’s side-by-side comparative visualization of chromosome X (*mus musculus*) as processed by the workflow. The browser has three different sections that promote interactive exploration of this complex data:

1. Selection Controls and Parameter Adjustments (Figure 3A). Scientists can select a feature (such as a gene), or a range from the sequence. If selections are made in either the structure view or the contact map view of the data, these fields are updated. Scientists can also adjust parameters such as the background color and the threshold for viewing the contact map.
2. Coordinated Structure and Contact Map View of Dataset (Figure 3B, Figure 3C). These panels contain coordinated views of data sets defined in the project.yaml. Users can select pieces of the sequence either in the structure view or the contact map view. Camera views of the data are synchronized across both visualizations.
1. 3. Track Views of Data Selections (Figure 3D). When a selection is made, the scientist can view detailed track information about that selection by selecting a variable to paint the structure with and then pressing the Created Data Track button. This will create a pairwise view of the track data, showing data from both visualizations at the same time. The entire set of tracks can be cleared by pressing the *Clear Data Tracks* button.

**Figure 3:**
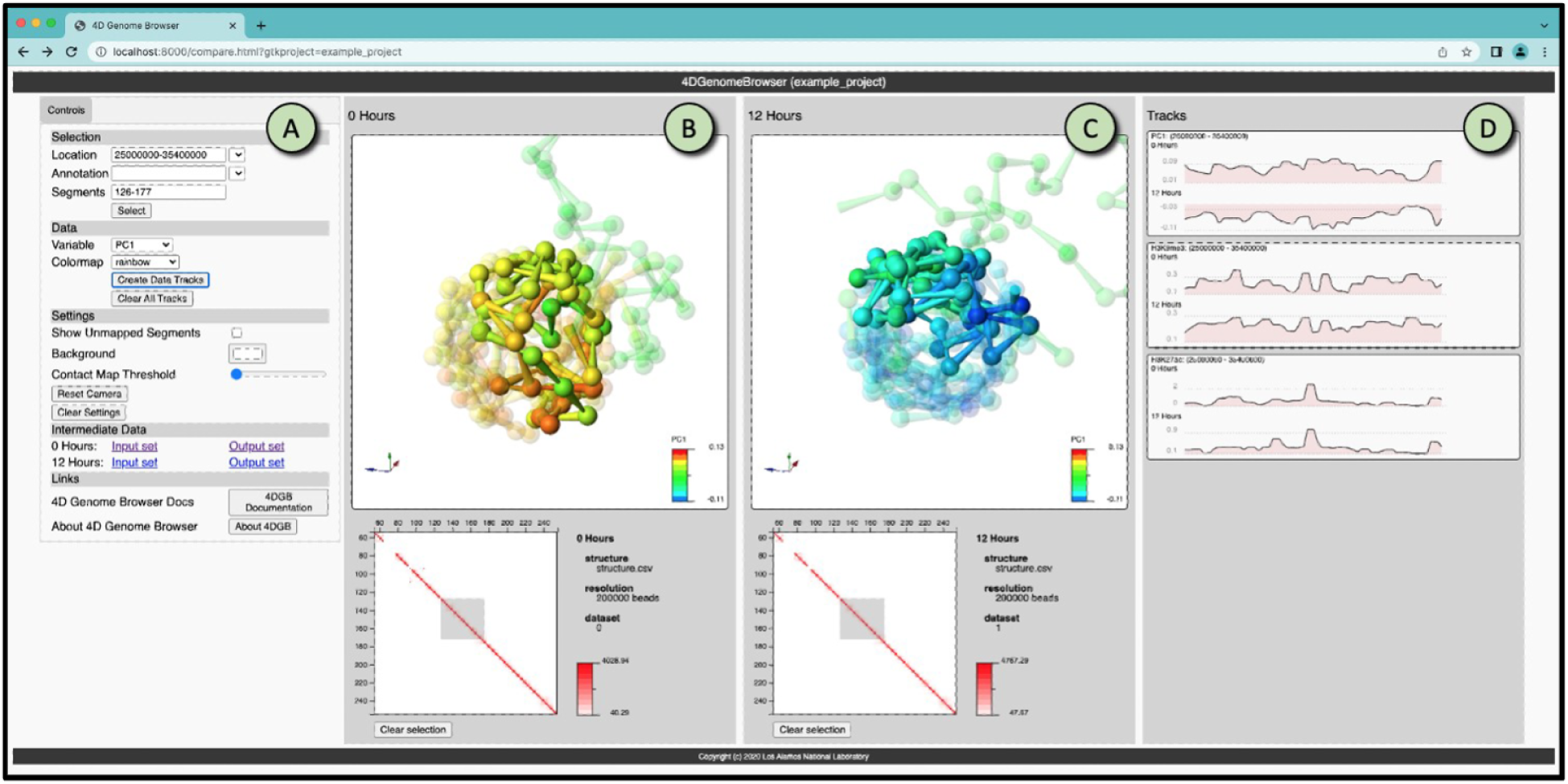
Screen capture of the 4D Genome Browser, showing the side-by-side comparison of two states of a chromosome (B and C) that are defined in the workflow. The browser consists of three main parts: the filter and view controls (A), the structure/contact map views (B, C), and the track inspector (D). A scientist uses the tool by selecting a portion of the sequence to highlight, and viewing tracks associated with that portion. The structure views rotate and scale in sync, and all selected, colored and highlighted data is synchronized between the views as well, allowing easy comparison of these complex datasets. The dataset shown is X chromosome (Mus musculus) previously described in [12, 30].

### Installing the Workflow

The following are prerequisites for running the workflow:

1. python v3.9 or higher must be installed on the system.
2. Docker must be installed on the system.

Once python and Docker are installed, the 4D Genome Browser Workflow can be installed by typing the following command in a shell:

pip install 4dgb-workflow

This will download and install a python package, making the tool accessible from the command line. It is left to the user to create a separate python environment for this tool if that is required.

### Running an Example Project

The 4DGB Workflow tool has command line options that make it easy to get started. The latest instructions for running the workflow can always be found in the online documentation. The following section summarizes those instructions. For simplicity of presentation, the commands shown in this section are specific to Linux and MacOS. Slight modifications to the commands are necessary on Windows. The online documentation has specifics.

The best path forward is to create a working directory specifically for this tool, and create and run the template project. The template project can then be edited to include your specific data.

To get started:

1. Install Docker, python and the 4dgb-workflow tool per the instructions in previous sections.
2. Create a new directory that will contain all data from your workflow runs. Multiple 4D Genome Browser Workflow projects can reside in this directory.
3. Open a shell, cd into your new directory, and run the following command. This will create and populate a new project directory with a default name. An optional –output argument can be added to define a specific name for the new directory.

### 4DGBWorkflow template

The new directory will contain the following data, which is used to run the project. These files show how to run the tool in various ways. Detailed instructions about them can be found in the online documentation.

./4DGB Project/

ENCLB571GEP.chr22.200kb.00hr.hic ENCLB571GEP.chr22.200kb.00hr.tracks.csv ENCLB870JCZ.chr22.200kb.12hr.hic ENCLB870JCZ.chr22.200kb.12hr.tracks.csv chr22.gff

features.csv project.full.yaml project.min-with-tracks.yaml project.min.yaml project.yaml

Next, run the workflow on the template data, using the command below. The tool will run, and report progress and errors to the shell.

### 4dgbworkflow run 4DGB Project

The first time you issue the run subcommand, the tool will download the Docker images needed to run the workflow. After the workflow is complete, a URL will be printed to the shell. Copy and paste that URL into a browser (Chrome, Safari and Firefox), and you should see the browser running on the data. If needed, a user may stop the workflow by typing ctrl+C in the shell. After the workflow has run, you can view the results at any time by running the view subcommand:

### 4dgbworkflow view 4DGB Project

This will start the server, and you can view your data in any browser using the URL displayed in the shell. After you have run the workflow to completion, the 3D structure data will not change, and it can be used over and over as you add or delete track data, bookmarks, annotations, etc. To do this, you edit your project.yaml file, then re-run the workflow. The workflow will determine how to create the data for the 4D Genome Browser without re-running the simulation. In this way, you can quickly view new data on the 3D structure.

Because the output from this workflow is a directory and files, results can be shared with other users, and across operating systems. The project directory can be copied or archived using the following shell commands, and uncompressed by anyone. The shared files will retain all process information, so the costly step of running the simulation will not have to be repeated. To copy an existing project:

cp -a existing project new project

To package up a project so that it can be moved and shared, preserving the file timestamps so that the simulation will not have to be run again:

tar -czvf myproject.tar.gz myproject

## Results and Discussion

Once the workflow is run, users can explore side-by-side coordinated views of the data (Figure 1). In this screen capture, the 4D Genome Browser displays analyzed Hi-C data from a recent experiment in the A549 cancer lung cell line [1]. In this study, A549 cells were treated with a histone deacetylase inhibitor (HDACi); samples were collected 24 and 48 hours post treatment and used to generate Hi-C sequencing data [1]. As an example, Figure 1 shows a view comparing the 3D renderings of control (left) and HDACi treated (right) Hi-C samples 24 hours post treatment. From these experiments, we observed increases in long-range chromatin contacts (i.e., interactions between linearly distal regions along chromosomes, greater than *>* 10^6^ base pairs apart) post HDACi treatment [1]. This effect was most prominent in distance decay profiles at 24 hours post treatment and while less pronounced, still visible 48 hours post treatment (Figure 5 in Venu et al., 2024 [1]). Supporting this original observation, increased connectivity along chromosome X between the p (red) and q (blue) arms of the chromosomes is observed, which are physically closer together in 3D space (Figure 1). Additionally, the p and q arms seem condensed in HDACi treated samples (compared to control) in 3D space post treatment (Figure 1). This visual comparison is typical of the kinds of insight possible with the 4D Genome Browser Workflow and 4D Genome Browser.

### Timing and Performance

It is useful to consider how large a problem can be effectively handled by this workflow. In practice, the most expensive computational step is the LAMMPS simulation, and the time this takes is proportional to the number of monomers that the simulation operates on. This in turn is related to both the size of the input chromosome and the resolution of that data. Multiple models of mouse chromosomes were generated on a 2020 MacBook Pro running MacOS, version 11.6.8 (Figure 4A). Though they differ in the number of Hi-C contacts, the build times remain consistent for all sample sizes of Hi-C contacts (Figure 4B). Thus, the build time does not appear to be impacted by the complexity of the input data (in particular, the complexity of the contact matrix).

**Figure 4:**
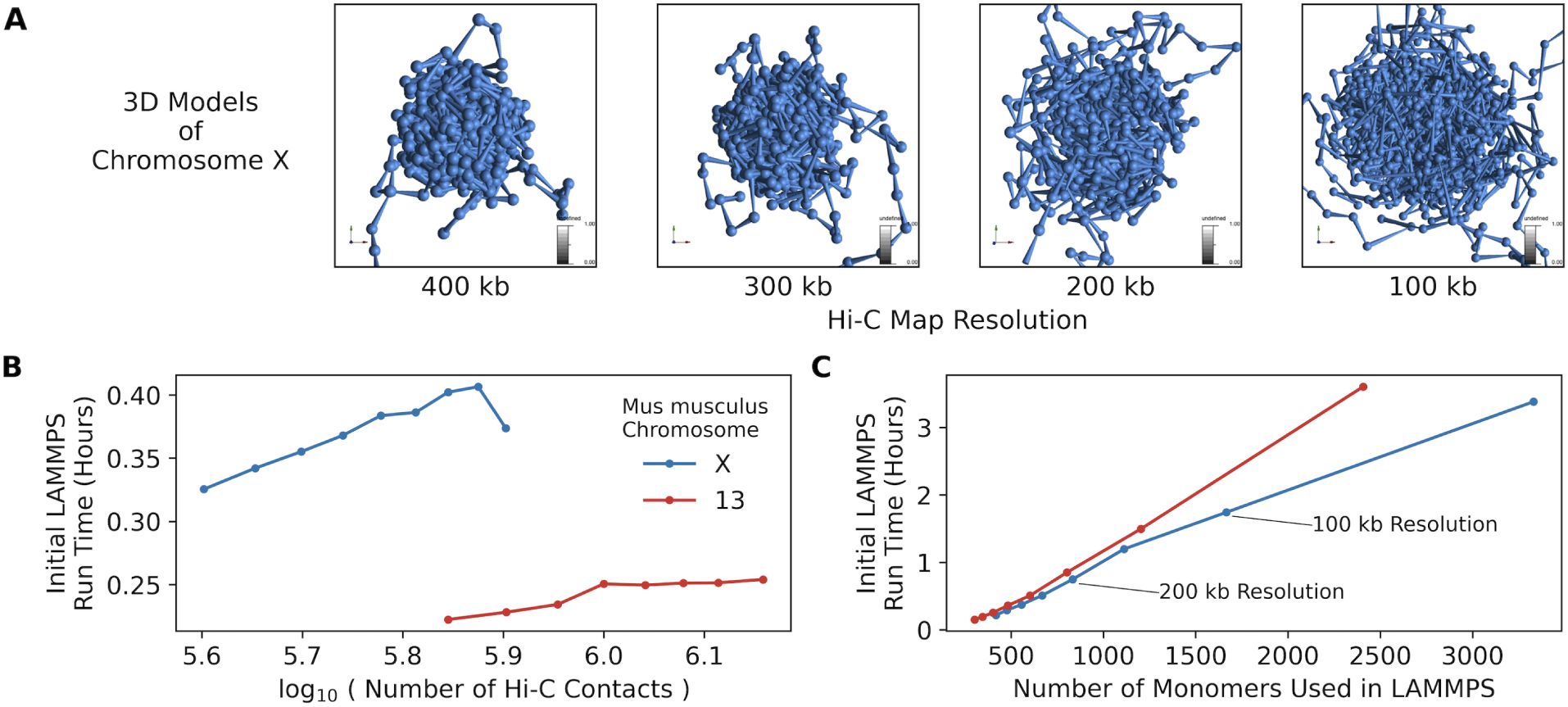
Performance information for the 4DGB workflow. A) Models of Mus musculus chromosome X at increasing genomic resolution (left to right, respectively). B) The LAMMPS build time (y-axis, minutes) as a function of number of Hi-C contacts (x-axis). Build time is not significantly affected by the overall number of paired end reads within a contact matrix (.hic). C) The LAMMPS build time (y-axis, minutes) as a function of the number of monomers set in LAMMPS (x-axis). Monomer count is determined by user defined resolution/bin size. In B and C, results using data along M. musculus chromosomes X and 13 are depicted in blue and orange respectively.

The build time increases linearly as a function of the number of monomers defined for the LAMMPS simulation (Figure 4C). As a practical matter, this is what will limit the usability of the tool, as LAMMPS simulations using over 3,000 monomers, which can model Hi-C data at 50 kb, takes approximately 200 minutes to complete. As stated before, once this build step is complete, additional data can be layered on the resulting model in the 4D Genome Browser without re-running the simulation(s).

### Integration of tool with workflow managers

Though the 4D Genome Browser Workflow is designed to work on a laptop, a containerized workflow has distinct advantages for web-based systems due to ease of installation, cross-platform performance, and lack of system dependencies. The tool has been integrated with the EDGE bioinformatics system [31], which allows users to upload data, initiate jobs, and view completed results through a simple graphical user interface. The 4DGB EDGE implementation utilizes the containerized workflow and executes it using data input from web forms and files uploaded by the user. This demonstrates the flexibility of the containerized workflow.

This instance of the 4DGB EDGE system is implemented on an internal network at Los Alamos National Laboratory. Authentication and data control are managed by the EDGE system. To use the system, the user logs into the EDGE system and is authenticated (Supplementary Figure S1), after which the user can browse a list of existing jobs, and inspect results from those jobs (Supplementary Figure S2). When a run is completed, the user can click on a link that brings up an instance of a new browser window running on the data requested. New jobs can be instigated by uploading new files, or using files that have already been uploaded to the system.

The integration with the EDGE framework is ongoing. In particular, each step in the workflow is being scaled up to handle higher resolution of both experimental and 3D structure data. Our goal is visualization of the entire human genome (∼3 billion base pairs); as such, the system requires a backend that can scale for exascale data. In addition, the visualization system must be able to perform selection, abstraction and visualization operations on billions of elements.

## Conclusions

Genome viewers have contributed to visualization, analysis, and discovery across a broad spectrum of genomic sciences for over a decade [32]. The Integrative Genome Browser and the UCSC Genome Browser are staples in the exploration of 1D genomic sequencing data [33,34]. As higher order data sets such as chromatin capture and Hi-C have become ubiquitous, researchers have developed several genomic viewers to visualize this type of data [35, 36]. A recent addition to the set of 3D genome browsers, MultiVis.js generates a reproducible workflow for iteratively exploring and tuning parameters for processing and visualizing 3D genomic data generated from Split-Pool Recognition of Interactions by Tag Extension experiments [37]. While these pipelines and applications are powerful tools, they do not support 3D modeling or 3D visualization of chromatin states.

The Nucleome Browser is a promising application that also furthers the visualization of 3D genomic data, supporting the classic visualizations of 1D sequencing tracks, integrated 2D heatmaps of Hi-C data, as well as 3D models of genomic and chromosome states [38]. This multimodel viewer incorporates and connects several databases and tools (such as the UCSC Genome Browser and HiGlass) to generate an integrated view of genomic data. While powerful, the 3D components of this tool are reliant on previously published or established nuclear models and data (such as those data produced by the NIH 4D Nucleome Consortium [39, 40]) and (to our knowledge) cannot create *de novo* 3D models of chromatin states generated from Hi-C data. The containerized code we have deployed here democratizes this complex process, and allows users to generate their own 3D models of chromosomes.

The 4DGB software package represents a unique toolset for transforming Hi-C datasets and functional genomic signal track information into 4D data that can be interactively explored in a web browser. The combination of the 4D Genome Browser Workflow and the 4D Genome Browser together provide an encapsulated, easily installed toolset that can be run on laptops, or integrated into web-based services. The comparative visualization application of the 4DGB Browser provides an opportunity for novel exploration of epigenomic datasets, helping scientists to understand interactions that occur in three dimensions across time (4D). It is our hope that this tool can help scientists better understand long-range interactions (across the entire genetic sequence) between genes and regulatory regions, as well as the mechanistic relationships between chromosome architecture and epigenetic modifications for deeper understanding of organismal function.

## Methods

### Data availability

Hi-C data from *Mus castaneus* and *Mus musculus* produced by Wang et al., 2018 [30] (and used in Lappala et al., 2021 [12]) used in timing experiments and figures are hosted on the Gene Expression Omnibus, with accession record GSE99991. The first replicate, with experimental accession record GSM2667262, was used here in testing.

Hi-C data from the ENCODE project were used in development and testing. Specifically the Hi-C data on the A549 cell line treated with dexamethasone (100 nM) at 0 and 12 hours were downloaded and processed. These are used in the example 4DGB Workflow project. The specific ENCODE accession names of data used are ENCLB571GEP and ENCLB870JCZ, for the 0 and 12 hour dexamethasone treated data, respectively.

Finally, a subset of Hi-C data from Venu et al., 2024 [1] was used in example visualizations. Specifically, the first replicate from control and HDACi (histone deacetylase inhibitor) treated A549 cells were used. These Hi-C samples are hosted on the Gene Expression Omnibus under accession record GSE282761.

### Data preparation and processing

Figure 1 and Figure 4 show examples of running the workflow to create and view 4D data of aligned sequencing records as featured in [12] and described previously [30] of the active and inactive mouse X chromosomes (Xa and Xi respectively) taken from the Gene Expression Omnibus, accession record: GSE99991. Specifically, the first replicate (experimental accession record: GSM2667262) of wild-type data for both two divergent strains of mice, *Mus castaneus* and *Mus musculus* (representing the Xa and Xi, respectively) was used in analysis. These records were converted from the .summary.txt.gz files to .hic files using custom python scripts. These scripts are described in the online documentation.

The 4D Genome Browser Workflow template project features Hi-C data from the ENCODE project [41], specifically a chromatin capture assay of A549 cancer lung cells, treated with dexamethasone (refer to online documentation for more details). The example data includes .hic files created by using HiCExplorer to align sequence reads and construct Hi-C maps [42]. First, the bwa-mem function was used to align sequenced reads and construct .sam files, separately for each read in pair, with default settings except for options A = 1, B = 4, E = 50, and L = 0 [43]. The paired, .sam files from above were used as input into the hicBuildMatrix function with a binsize of 200 kb. The hicFindRestSites was used to find restriction cut-sites for the MboI enzyme. The Hi-C map construction (in .h5 format) was limited to chromosomes 22.

For Hi-C data from Venu et al., 2024 [1], raw data was processed with the Juicer processing pipeline with default settings [27]. Details on DNA library construction and sequencing are previously published and listed elsewhere, please see [1, 11].

## Availability and Requirements

Project name: 4DGBWorkflow

Project home page: https://github.com/4DGB/4DGBWorkflow Associated Repository: https://github.com/4dgb

Additional tools for file conversion: https://github.com/4DGB/hic-converter Online Documentation: https://4dgb.readthedocs.io

Operating system(s): Linux (dev/build); MacOS, Linux, Windows (tool deployment) Programming language: Python, javascript

Other requirements: Docker desktop License: BSD three clause

Any restrictions to use by non-academics: None

## Declarations

### Acknowledgements

The authors would like to thank the members of the Lee Lab, Steadman Lab, and members of the Genomics and Bioanalytics Group at Los Alamos National Lab for beta tests and feature suggestions for the 4D Genome Browser. We would also like to thank the lab of Tim Reddy of Duke University for producing data used here from the ENCODE consortium.

### Contributions

DHR, CR, CRS, and SS wrote the paper. AL developed underlying code, data, and methods for running 4DHiC pipeline and LAMMPS simulations to produce 3D chromosome models. DHR and CR generated figures. CR performed timing experiments. CT developed code underlying the 4DGB workflow and tool. CRS, KS, JTL and SS provided funding.

### Ethics approval and consent to participate

Not applicable.

### Consent for publication

Not applicable.

## Supplementary Materials

**Figure S1:**
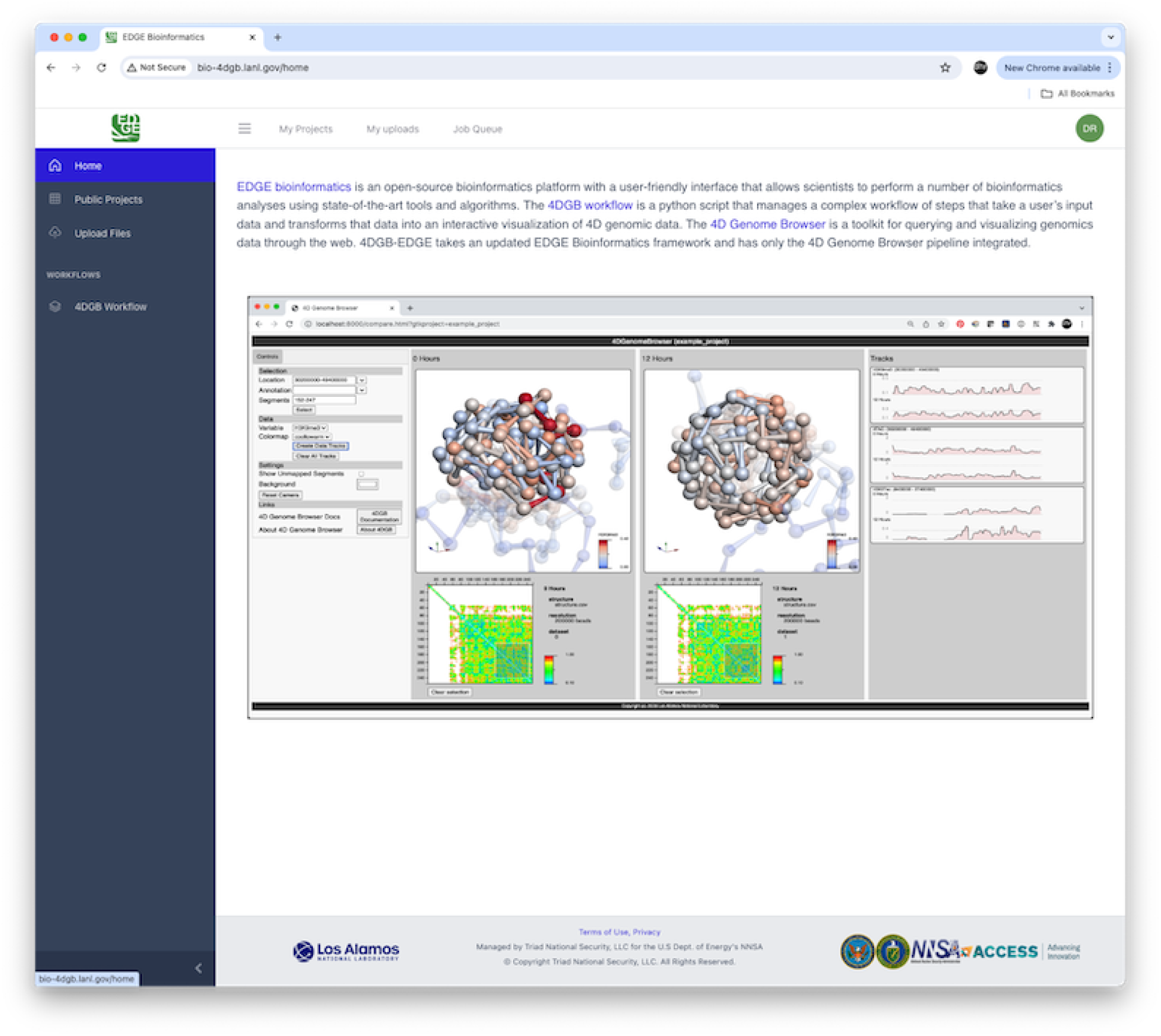
4DGB on EDGE Screen capture of the 4DGB Edge home screen, showing a capture of the tool instructions and both side and top menus for access to web-based capabilities.

**Figure S2:**
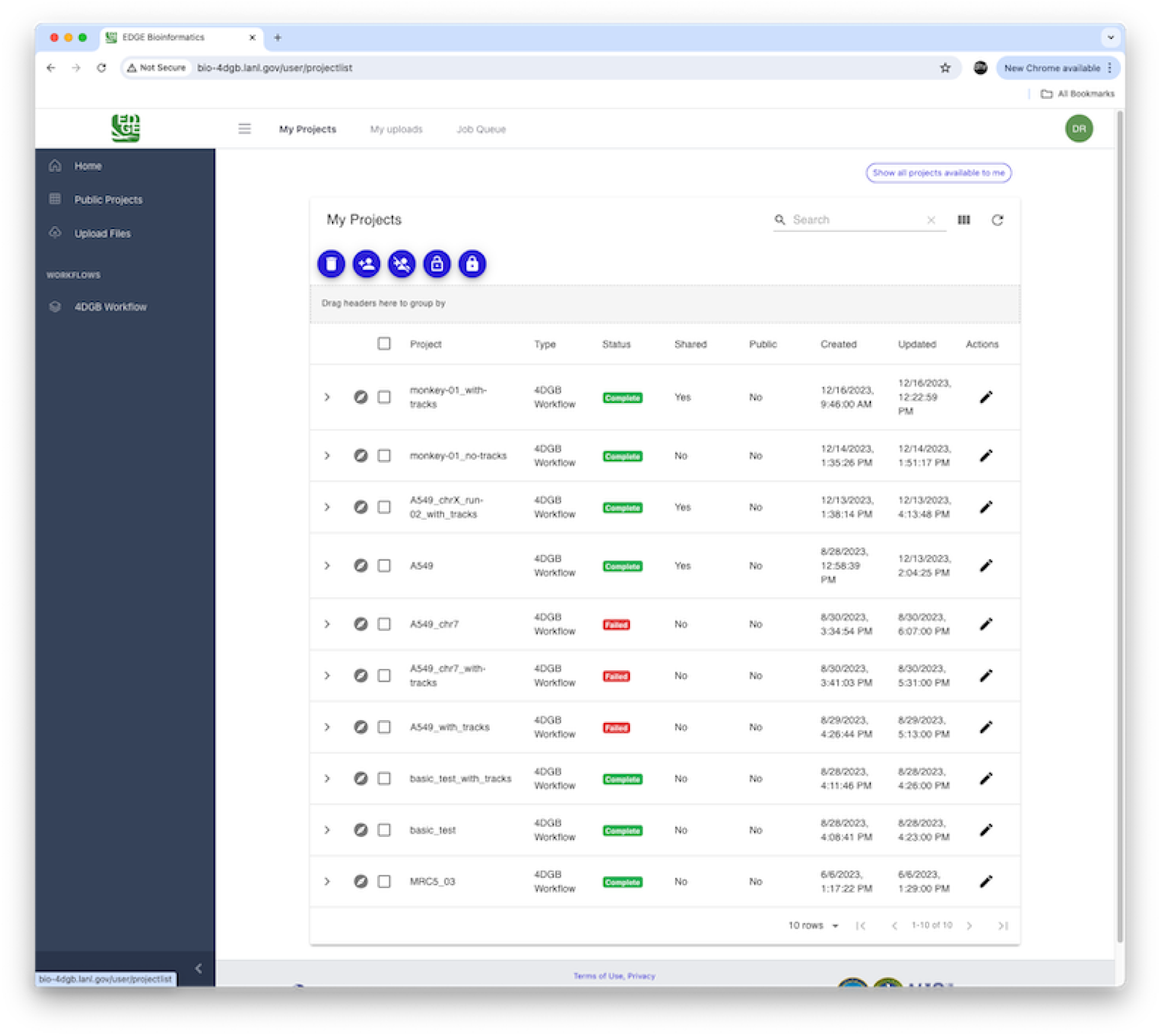
4DGB on EDGE Screen capture of the 4DGB EDGE implementation, showing the list of existing jobs, which the user can navigate to view the final data products.

