## Supplemental Figure 1 for "From 2D to 4D: a Containerized Workflow and Browser to Explore Dynamic Chromatin Architecture"

EDGE Bioinformatics

Not Secure bio-4dgb.lanl.gov/home

New Chrome available

All Bookmarks

My Projects My uploads Job Queue

DR

- Home
- Public Projects
- Upload Files

WORKFLOWS

- 4DGB Workflow

EDGE bioinformatics is an open-source bioinformatics platform with a user-friendly interface that allows scientists to perform a number of bioinformatics analyses using state-of-the-art tools and algorithms. The 4DGB workflow is a python script that manages a complex workflow of steps that take a user's input data and transforms that data into an interactive visualization of 4D genomic data. The 4D Genome Browser is a toolkit for querying and visualizing genomics data through the web. 4DGB-EDGE takes an updated EDGE Bioinformatics framework and has only the 4D Genome Browser pipeline integrated.

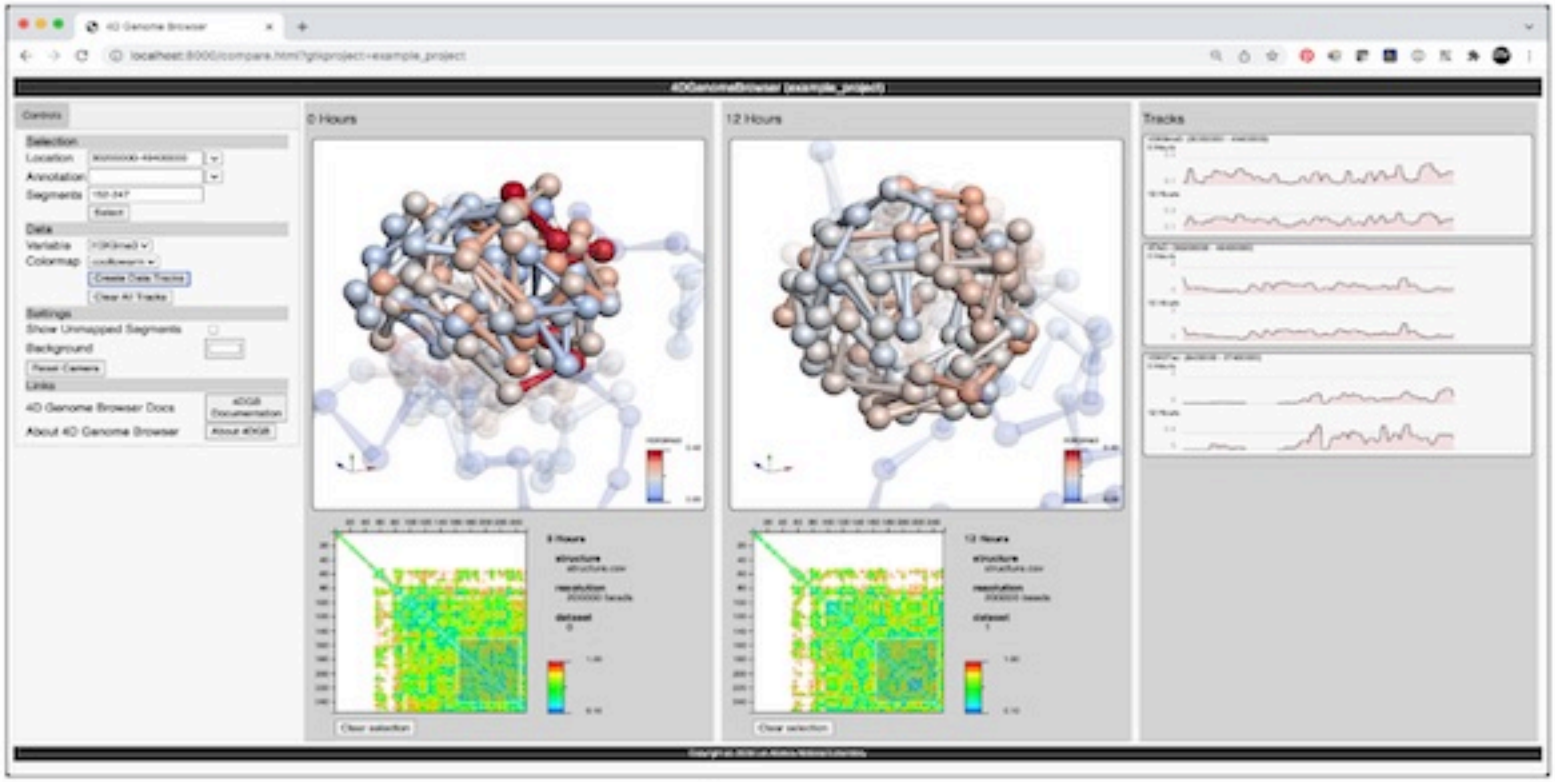


Los Alamos NATIONAL LABORATORY

Terms of Use, Privacy

Managed by Triad National Security, LLC for the U.S Dept. of Energy's NNSA

© Copyright Triad National Security, LLC. All Rights Reserved.

NNSA ACCESS Advancing Innovation

bio-4dgb.lanl.gov/home
