## Supplementary figures and images for "From 2D to 4D: a Containerized Workflow and Browser to Explore Dynamic Chromatin Architecture"

### Supplemental Figure 2

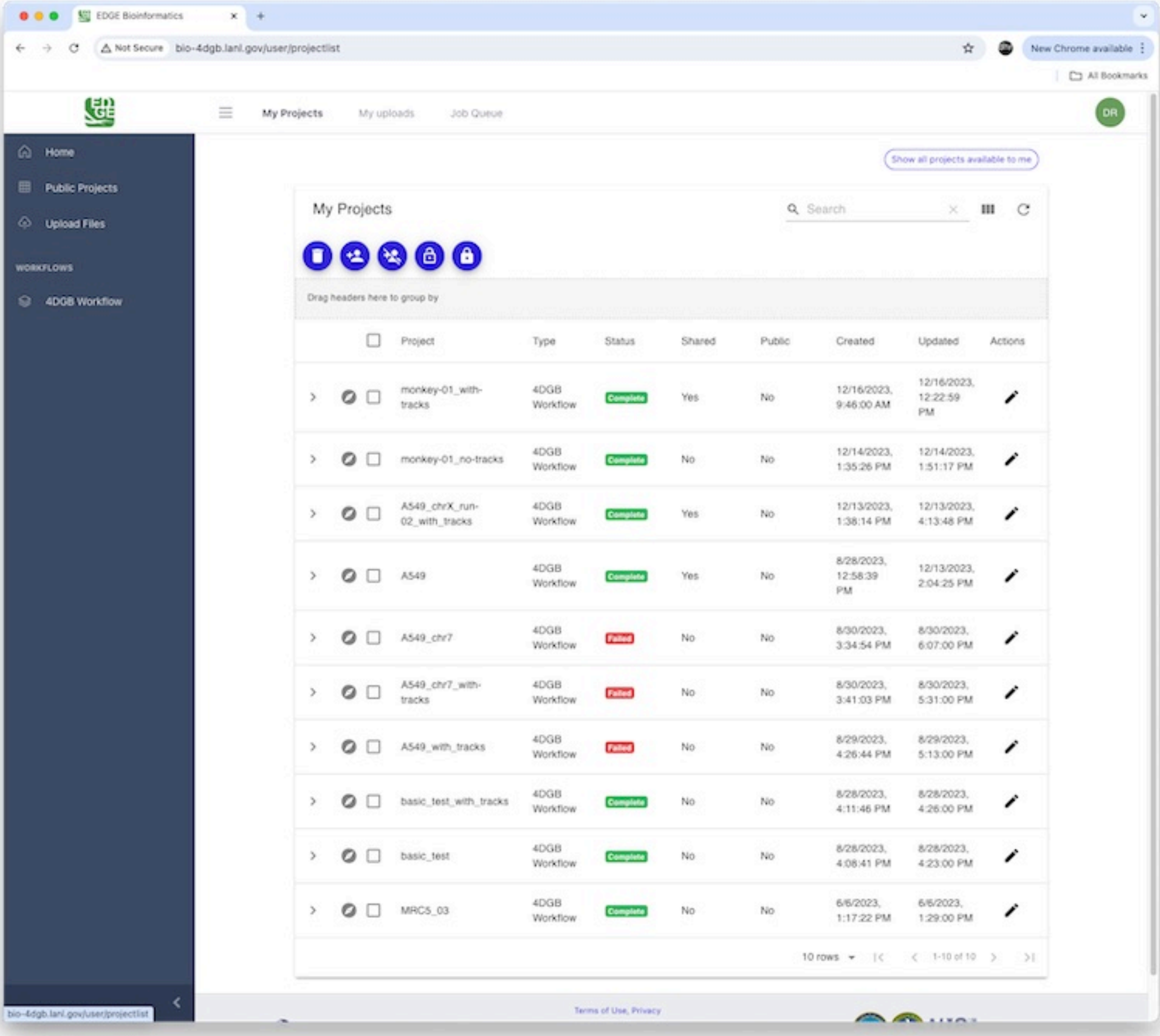
